# Seroepidemiological survey on pigs and cattle for novel K88 (F4)-like colonization factor detected in human enterotoxigenic *Escherichia coli*

**DOI:** 10.1101/2021.05.01.442292

**Authors:** Yoshihiko Tanimoto, Miyoko Inoue, Kana Komatsu, Atsuyuki Odani, Takayuki Wada, Eriko Kage-Nakadai, Yoshikazu Nishikawa

**Author notes:** **Author for correspondence** Yoshikazu Nishikawa, Faculty of Human Sciences; Tezukayamagakuin University; Osaka, Japan.

## Abstract

Enterotoxigenic *Escherichia coli* (ETEC) strains that express various fimbrial or nonfimbrial colonization factors and enterotoxins are critical causes of diarrheal diseases. Human ETEC serotype O169:H41 (O169) has been a representative of epidemic ETEC worldwide; the organism shows massive adherence to HEp-2 cells similar to enteroaggregative *E. coli*. Previously, we determined the complete sequence of the unstable virulence plasmid, pEntYN10. The plasmid included a unique set of genes encoding a novel colonization factor (CF) resembling K88 (F4) of porcine ETEC, in addition to CS6, a well-known representative CF of human ETEC, and another novel CF similar to CS8 (CFA/III) of human ETEC. To determine whether the K88-like CF (after this, K88_O169_) allows the organisms to infect domestic animals like the original K88-harboring strains that can cause diarrhea in piglets, samples were tested for antibodies against recombinant proteins of possible paralogous adhesins, FaeG1 and FaeG2, from K88_O169_ and the FaeG of typical K88 (F4). The seroepidemiological study using recombinant antigens (two paralogs FaeG1 and FaeG2 from K88_O169_) showed reactivity of porcine (18.0%) and bovine (17.1%) sera to K88_O169_ FaeG1 and/or FaeG2 antigens on indirect ELISA tests. These results suggest that *E. coli* with K88_O169_ adhesin can infect various hosts, including pigs and cattle. This is the first report of domestic animals having antibodies to K88_O169_ of human ETEC. Although human ETEC had been thought to be distinguished from those of domestic animals based on colonization factors, zoonotic strains may conceal themselves among human ETEC organisms. The concept of One Health should be adopted to intervene in ETEC infections among animals and humans.

## Introduction

Enterotoxigenic *Escherichia coli* (ETEC) is a diarrheagenic *E. coli* that causes diarrhea not only in humans but in various animals, such as pigs, cattle, and sheep [1]. Regardless of host, colonization of the intestinal epithelia is an essential first step in the pathogenesis of ETEC infection mediated by a variety of colonization factors (CFs) [2]. Subsequently, ETEC secretes host-damaging toxins such as heat-labile (LT) and/or heat-stable (ST) enterotoxins, which give rise to intestinal symptoms such as diarrhea [3,4].

ST-producing ETEC O169:H41 (O169) was first identified as an etiological serotype in human foodborne cases in Japan [5]. Previously, we reported that the O169 strain YN10 harbors the unstable large plasmid pEntYN10, which codes for genes of three CFs, CS6 and two novel CFs called resembling CS8 (so-called CFA/III) and F4 (so-called K88). The operon coding the novel K88-like CF (hereinafter, K88_O169_) has two paralogous major adhesin-like subunits, *faeG1* and *faeG2*, which have 37%-44% amino acid homology with *faeG* of the original K88 [6].

K88 and its related CFs have been found with bacteria isolated from a variety of hosts. K88 was initially identified as a CF of swine ETEC [7,8], whereas CS31A was reported as a K88-related CF from bovine ETEC [9]. Thus, *E. coli* strains with analogous CFs were isolated from multiple hosts, suggesting that K88-related CFs can lead to colonization in various host species. Furthermore, an ETEC/Shiga toxin-producing *E. coli* (STEC) hybrid strain was recently isolated from a patient with Hemolytic-Uremic Syndrome (HUS); the organism possessed an F4-like adhesin of the protein sequence that was similar to CS31A of bovine ETEC [10]. The presence of multiple CFs in O169 may indicate the potential for multi-host transmission: a complex zoonotic network may be supported, and this contradicts the one-to-one host-pathogen relationship that has been commonly accepted for ETEC.

This study examined the seroepidemiological status of livestock against K88_O169_ as a preliminary investigation to test this hypothesis. Antibody titers against K88_O169_ of pigs and cattle farmed in various areas in Japan were examined by an indirect ELISA method using recombinant proteins of the adhesins, FaeG.

## Materials and Methods

### Sample collection

In this study, sera of 200 pigs and 105 cattle collected from June 2019 to March 2020 was kindly provided by the Osaka City Meat Hygiene Inspection Center for serological assays. This facility is partnered with the slaughterhouse and is responsible for hygiene inspections of meat supplied to the metropolitan area and also conducts research on livestock animals transported from a wide range of regions in Japan. Breeders and the prefectures where the animals were raised are listed in **Supporting Table 1**.

### Recombinant protein

Recombinant proteins were created using the pET expression system (Novagen). The codon-optimized typical *faeG* gene of the original K88 (K88ac variant, accession ID: AJ616239) [11] was synthesized by Eurofins Genomics KK (Tokyo, Japan). *faeG1* and *faeG2* genes of K88_O169_ were amplified by PCR with O169 plasmid pEntYN10 [6] as the template. The primers are described in **Supporting Table 2**. The primers were designed to remove signal peptides predicted using SignalP 5.0 web-based resource [12] from the recombinant proteins. These gene products and pET30a vector (Novagen) were digested with *Sal*I and *Not*I restriction enzymes and ligated. The resulting plasmids (pET30a-*faeG*, pET30a-*faeG1,* and pET30a-*faeG2*) were recovered by transformation into *E. coli* BL21 (DE3) with selection for kanamycin resistance. After the bacteria were grown at 37°C for 3 h, protein expression was induced with 1 mM of isopropyl-β-D-thiogalactopyranoside (IPTG; Takara Bio) at the final concentration at 25°C. After overnight incubation, the cells were solubilized with 8 M urea. The recombinant protein was purified using the His60 Ni gravity column purification kit (Takara Bio) according to the manual instruction. Finally, the solution was exchanged with phosphate buffered saline (PBS) using Amicon Ultra-15 10K Centrifugal Filter Devices (Merck). Purity was confirmed by the absence of contamination bands on SDS-PAGE.

### Antiserum

Antiserum to the original K88 was purchased from Denka Seiken (*E. coli* K88 rabbit antiserum). Rabbit polyclonal antibodies against FaeG1 and FaeG2 developed by Eurofins Genomics Antibody Service were used as positive controls.

### Indirect ELISA

Sample sera were examined for the antibodies against FaeG of K88, FaeG1 and FaeG2 of K88_O169_ with indirect ELISA methods. Purified protein was coated on 96-well half-area plates (Corning) at 0.1 μg/mL with coating buffer (13 mM Na_2_CO_3_, 35 mM NaHCO_3_) overnight 4°C. After washing three times with PBS-T (PBS containing 0.05% Tween-20), plates were blocked by adding blocking buffer (0.4% Block Ace, Dainippon Pharmaceutical) and incubated for 1-2 h. After three washes, sample sera (1:200 dilution in PBS-T) and antibodies (1:1000 dilution) as positive control were added to the plate and incubated for 1 h. After three washes, a 1:30,000 dilution of the second antibody solution was added to each well and incubated for 1 h. The second antibodies used for ELISA were as follows: HRP-linked anti-whole rabbit IgG donkey serum (GE Healthcare, #NA934) for detection of the positive control, HRP-linked anti-pig whole IgG goat antibodies (abcam, #ab6915) for porcine samples, and HRP-linked anti-cow whole IgG goat antibodies (abcam, #ab102154) for bovine samples. After three washes, TMB substrate (TMB Microwell Peroxidase Substrate System, KPL) was added to the plates, incubated for 5 min in the dark, stopped with 1 M H_2_SO_4,_ and measured at wavelengths of 450 nm/550 nm with a microplate reader (Wallac 1420 ARVOsx, Perkin Elmer). Sample to positive (S/P) ratio (%) was calculated as follows: (Sample serum – blank) / (Positive control antibody – blank). S/P ratio > 0.4 was considered as positive [13,14].

### Statistical analysis

The results were statistically analyzed using Prism software (GraphPad ver. 8.4.3). Fisher’s exact test was employed to assess the association between porcine and bovine seroprevalence.

## Results

Anti-FaeG1 and -FaeG2 antibodies in porcine and bovine sera were measured to consider whether K88_O169_ can be another option for ETEC. Among the samples tested, a portion of the porcine (18.0%, 36/200) and bovine (17.1%, 18/105) sera reacted to one or both of FaeG1 and FaeG2 antigens of K88_O169_ (**Table 1, Figs. 1, 2**). The prevalence of K88_O169_ antibodies reacting to FaeG1 and/or FaeG2 did not show any statistical difference between porcine and bovine sera. Antibodies against FaeG1 were detected among 16.0% (32/200) of pigs and 12.4% (13/105) of cattle, respectively; anti-FaeG2 antibodies were found at 7.5% (15/200) of pigs and 9.5% (10/105) of cattle. Prevalence of antibodies to each of FaeG1 and G2 showed no statistical difference between porcine and bovine sera (FaeG1; P=0.4979, FaeG2; P>0.5199). Anti-FaeG1 single positive sera were more among cattle (n=8, 7.6%) than pigs (n=9, 4.5%) (P=0.2968) (**Table 2**). Anti-FaeG2 single positive sera were significantly higher among cattle (n=4, 3.8%; P=0.0495) than pigs (n=1, 0.5%).

**Table 1.** Prevalence of antibodies against FaeG1 and FaeG2 among porcine and bovine sera

| Antigens (K88<sub>O169</sub>) |  | Porcine sera (n=200) | Bovine sera (n=105) | P value† |
| --- | --- | --- | --- | --- |
| FaeG1 | FaeG2 | (%) | (%) |  |
| + | + | 11 | 5 | P>0.9999 |
|  |  | (5.5%) | (4.76%) |  |
| + | - | 21 | 8 | P=0.5386 |
|  |  | (10.5%) | (7.62%) |  |
| - | + | 4 | 5 | P=0.2834 |
|  |  | (2.0%) | (4.76%) |  |
| - | - | 164 | 87 | P>0.9999 |
|  |  | (82.0%) | (82.9%) |  |

†Fisher's exact test was employed to assess the association between porcine and bovine seroprevalence.

**Fig. 1.**
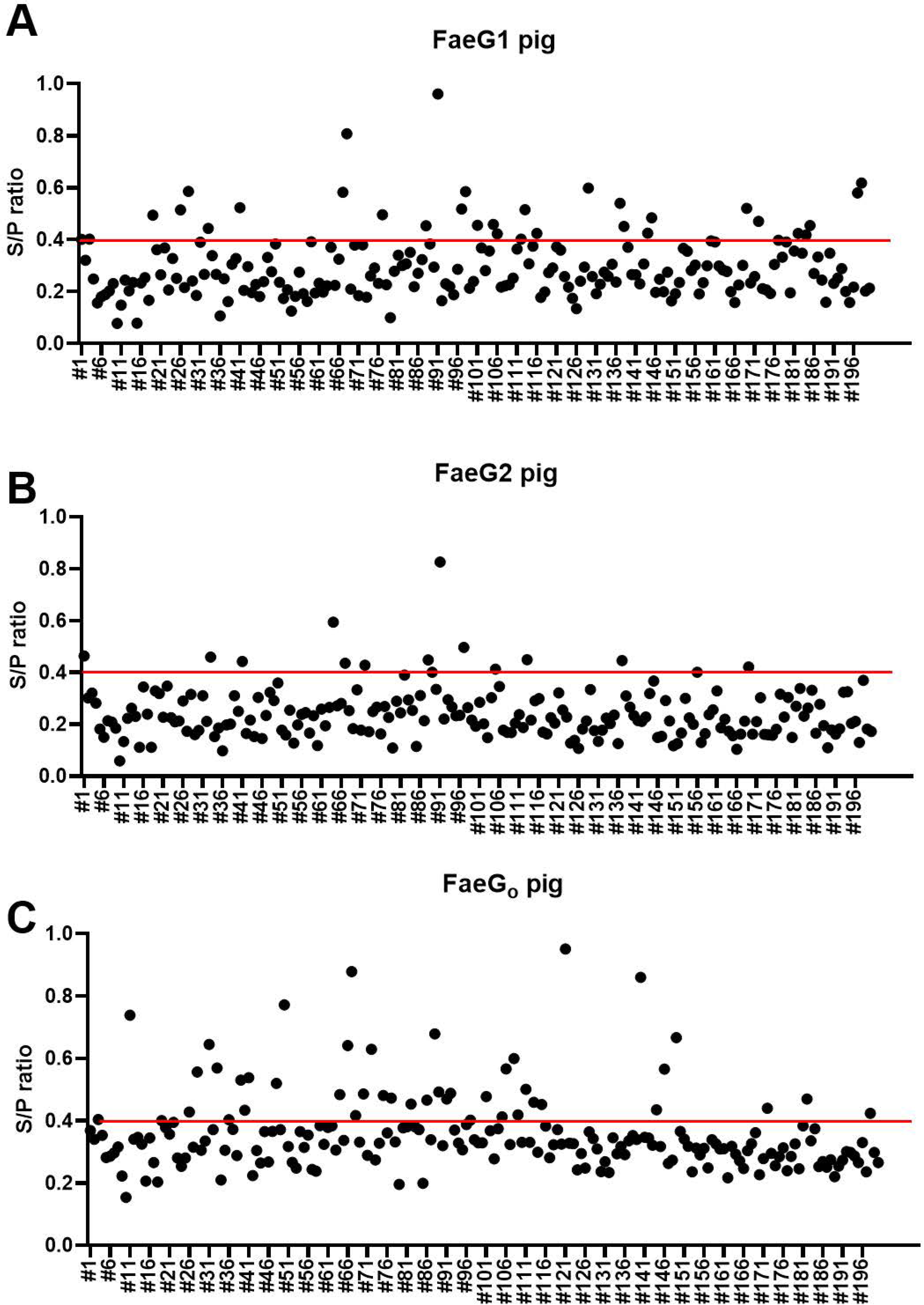
Indirect ELISA plot of porcine serum. Individual porcine sera against FaeG1 (A), FaeG2 (B), and FaeG_O_ (C) plotted as the S/P ratio.

**Fig. 2.**
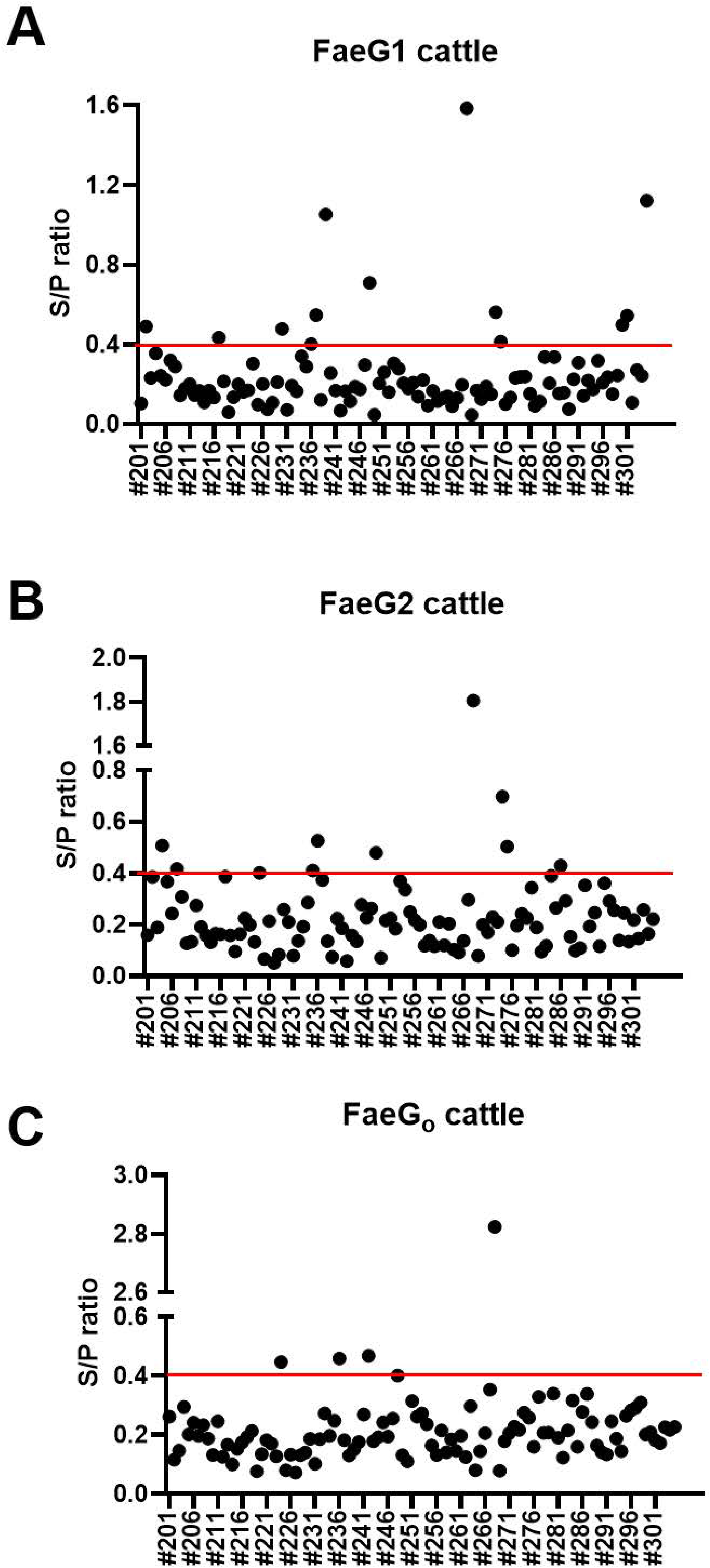
Indirect ELISA plot of bovine serum. Individual bovine sera against FaeG1 (A), FaeG2 (B), and FaeG_O_ (C) plotted as the S/P ratio.

**Table 2.** Prevalence of antibodies against FaeGo, FaeG1, and FaeG2 among porcine and bovine sera

| Antigens |  |  | Porcine sera<br>(n=200)<br>No. of samples (%) | Bovine sera (n=105)<br>No. of samples (%) | P value† |
| --- | --- | --- | --- | --- | --- |
| FaeGo (K88) | FaeG1(K88<sub>O169</sub>) | FaeG2 (K88<sub>O169</sub>) |  |  |  |
| + | + | + | 8 (4.0) | 3 (2.9) | P=0.7535 |
| + | + | - | 12 (6.0) | 0 (0) | **P=0.0098 |
| + | - | + | 3 (1.5) | 1 (1.0) | P>0.9999 |
| - | + | + | 3 (1.5) | 2 (1.9) | P>0.9999 |
| + | - | - | 21 (10.5) | 1 (1.0) | **P=0.0017 |
| - | + | - | 9 (4.5) | 8 (7.6) | P=0.2968 |
| - | - | + | 1 (0.5) | 4 (3.8) | *P=0.0495 |
| - | - | - | 143 (71.5) | 86 (81.9) | P=0.0515 |

†Fisher's exact test was employed to assess the association between porcine and bovine seroprevalence.

In contrast, when we evaluated the prevalence of antibodies against FaeG of K88ac (hereinafter, FaeG_O_, to reflect its originality), a typical adhesion factor of porcine ETEC, the antibodies to FaeG_O_ were obviously more prevalent among pigs than cattle (**Table 3, Figs. 1, 2**).

**Table 3.** Prevalence of antibodies against FaeGO among porcine and bovine sera

| Antigens (K88) | Porcine sera (n=200) | Bovine sera (n=105) | P value† |
| --- | --- | --- | --- |
| FaeGo | (%) | (%) |  |
| + | 44<br>(22.0%) | 5<br>(4.8%) | ***P<0.0001 |
| - | 156<br>(78.0%) | 100<br>(95.2%) | ***P<0.0001 |

†Fisher's exact test was employed to assess the association between porcine and bovine seroprevalence.

Comparison of epitopes of FaeG_O_, FaeG1, and FaeG2 adhesins shows apparent differences in amino acid sequences (**Fig. 3**) [15]. Individuals positive for antibodies to FaeG_O_ of K88 did not necessarily cross-react to FaeG1 or FaeG2 of K88_O169_. A total of 27 samples showed a positive reaction to Fae G1 or FaeG2 antigens but not FaeG_O_, and 22 anti-FaeG_O_ positive samples were negative to FaeG1 or FaeG2 antigens (**Table 2**).

**Fig. 3.**
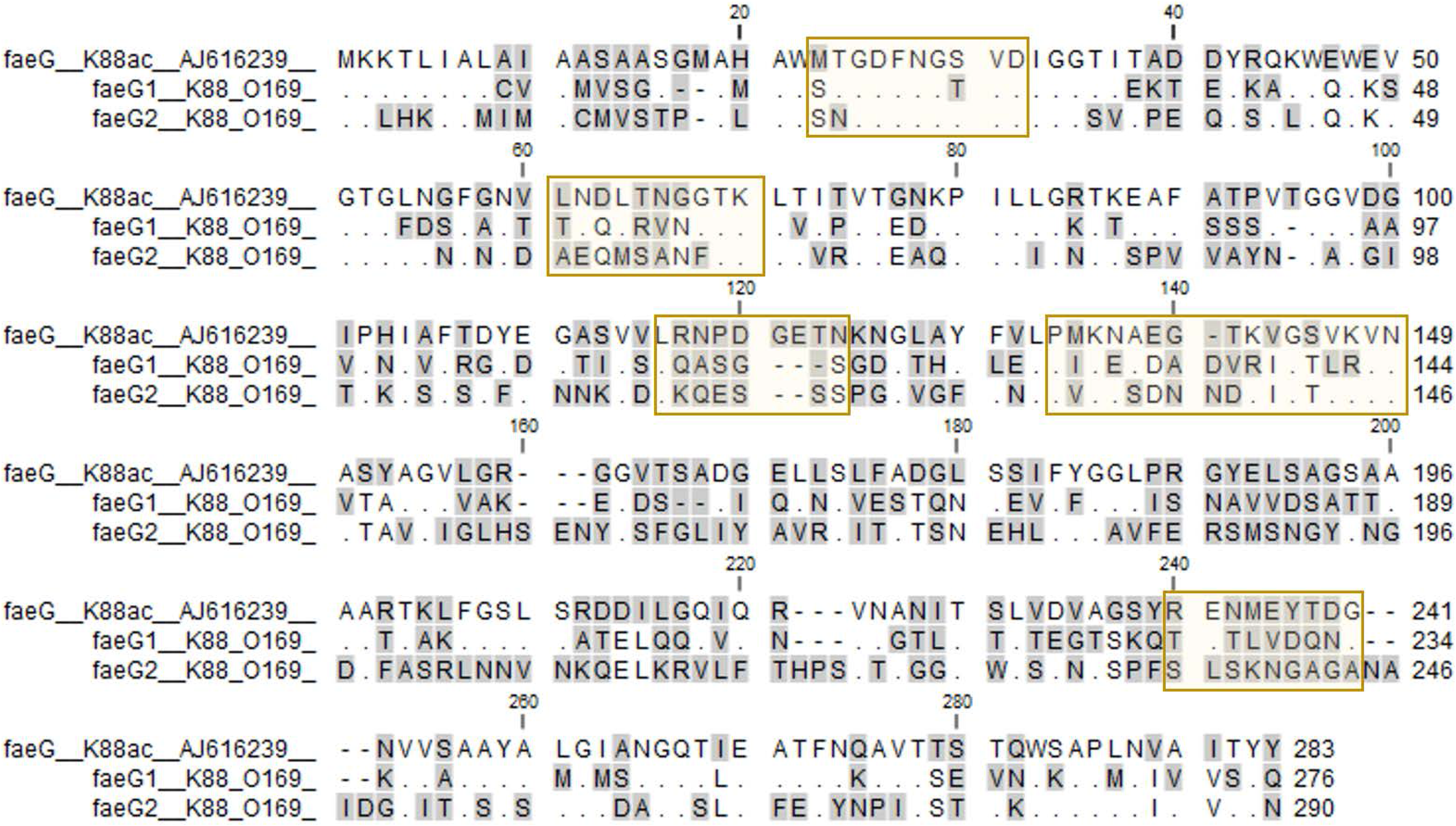
Epitope comparison of FaeG_o_, FaeG1 and FaeG2. The amino acid sequences of FaeG_O_, FaeG1, and FaeG2 were aligned using ClustalW. Residues identical to FaeG_O_ are shown as dots. Hyphens indicate positions where residues are missing. The residues enclosed in a square are the epitope referenced in a previous study [15].

## Discussion

The O169 plasmid pEntYN10 has a total of three CFs (CS6, K88-like, and CFA/III-like) [6]. CS6 is recognized as one of the representative CFs for humans; however, the role of the other two CFs remained to be elucidated. Since the O169 plasmid can be easily eliminated from the bacteria in vitro [6], multiple CFs found in the plasmid evoked us the possibility that they make the bacteria infect various species of hosts consecutively. We evaluated seroepidemiologically if ETEC possessing K88_O169_ could infect pigs because K88 is a classical colonization factor of porcine ETEC.

The porcine and bovine sera reacted to one or both of FaeG1 and FaeG2 antigens of K88_O169_. These results suggest that K88_O169_-positive *E. coli* infect pigs and cattle. Further, the results were not biased toward any particular region because the sera samples were collected from animals reared in various prefectures (**Supporting Table 1**). On the other hand, the FaeG_O_ positivity rate was significantly higher among pigs than cattle (P<0.0001), which seroepidemiologically reflects that ETEC possessing K88 colonize more prevalently among pigs. This is consistent with previous reports that the original K88 is a typical adhesion factor of ETEC that causes diarrhea in piglets [16,17].

We considered the possibility that anti-FaeG_O_ cross-reacts with FaeG1 and FaeG2. However, epitopes of FaeGo, FaeG1, and FaeG2 adhesins show differences in amino acid sequences. Individuals positive for antibodies to FaeGo, FaeG1, or FaeG2 alone were observed in both pigs and cattle. These results suggest that individuals infected with the bacteria harboring FaeG1 and FaeG2 are prevalent in these species; the positive reactions to these novel antigens are unlikely due to the cross-reaction of antibodies to FaeG_O_.

CF is generally recognized as being the molecule that determines host specificity. However, our data showed that anti-K88_O169_ was prevalent among bovine sera and porcine sera, suggesting that K88_O169_ may provide the strain with a mechanism that allows it to infect multiple hosts. The presence of CS8-like CF plasmid, the third and novel one of the O169 strain, may also enhance survival strategies by expanding the host spectrum bacteria in which it is present. Interestingly, the virulence plasmid pEntYN10 tended to be quickly eliminated *in vitro* because there are fewer genes associated with plasmid maintenance [6]. This plasmid is maintained in ETEC O169 despite this disadvantage, suggesting that the plasmid may provide its bacterial hosts with high fitness to survive in the bacteria.

In conclusion, antibodies to K88_O169_ antigens are prevalent among pigs and cattle. Although specific hosts where the K88_O169_ acts as CF remain to be revealed, O169 organisms may expand their niches through a unique and complex repertory of CFs. Not only O169 but other serogroups of ETEC possessing K88_O169_ can be transmitted from domestic animals to humans or from humans to domestic animals using different adhesins across hosts. We should pay attention to ETEC as a possible pathogen of zoonosis.

## Acknowledgements

The authors thank veterinarians Mitsuhiro Tsujimoto, Tomofumi Maehara, Norihide Kuriyama, Hideki Oshima, and Yusuke Kataoka, at the Osaka City Meat Hygiene Inspection Center, for providing porcine and bovine sera.

## Author’s contributions

YT and YN designed the study. YT, MI, KK, and AO carried out the experiments. YT analyzed data. EKN was involved in planning and supervised the work. YT, TW, and YN wrote the paper. All authors received the final manuscript version for approval.

