## Supplemental Tables for "Seroepidemiological survey on pigs and cattle for novel K88 (F4)-like colonization factor detected in human enterotoxigenic *Escherichia coli*"

Supporting Table 1. Swine sera used in this study

| No. | Collection date | Vendor | Prefecture |
|---|---|---|---|
| 1 | 2019/6/17 | A | Nara |
| 2 | 2019/6/17 | A | Nara |
| 3 | 2019/6/17 | A | Nara |
| 4 | 2019/6/17 | A | Nara |
| 5 | 2019/6/17 | A | Nara |
| 6 | 2019/6/17 | B | Mie |
| 7 | 2019/6/17 | B | Mie |
| 8 | 2019/6/17 | B | Mie |
| 9 | 2019/6/17 | B | Mie |
| 10 | 2019/6/17 | B | Mie |
| 11 | 2019/6/17 | C | Hyogo |
| 12 | 2019/6/17 | C | Hyogo |
| 13 | 2019/6/17 | C | Hyogo |
| 14 | 2019/6/17 | D | Hiroshima |
| 15 | 2019/6/17 | D | Hiroshima |
| 16 | 2019/6/18 | E | Wakayama |
| 17 | 2019/6/18 | E | Wakayama |
| 18 | 2019/6/18 | E | Wakayama |
| 19 | 2019/6/18 | F | Mie |
| 20 | 2019/6/18 | F | Mie |
| 21 | 2019/6/18 | F | Mie |
| 22 | 2019/6/18 | F | Mie |
| 23 | 2019/6/18 | F | Mie |
| 24 | 2019/6/18 | G | Hiroshima |
| 25 | 2019/6/18 | G | Hiroshima |
| 26 | 2019/6/18 | G | Hiroshima |
| 27 | 2019/6/18 | G | Hiroshima |
| 28 | 2019/6/18 | G | Hiroshima |
| 29 | 2019/6/18 | H | Mie |
| 30 | 2019/6/18 | H | Mie |
| 31 | 2019/6/20 | I | Hyogo |
| 32 | 2019/6/20 | I | Hyogo |
| 33 | 2019/6/20 | I | Hyogo |
| 34 | 2019/6/20 | I | Hyogo |
| 35 | 2019/6/20 | I | Hyogo |
| 36 | 2019/6/20 | I | Hyogo |
| 37 | 2019/6/20 | J | Ehime |
| 38 | 2019/6/20 | E | Wakayama |
| 39 | 2019/6/20 | E | Wakayama |
| 40 | 2019/6/20 | J | Ehime |
| 41 | 2019/6/20 | J | Ehime |
| 42 | 2019/6/21 | K | Nara |
| 43 | 2019/6/21 | K | Nara |
| 44 | 2019/6/21 | K | Nara |
| 45 | 2019/6/21 | L | Okayama |
| 46 | 2019/6/21 | L | Okayama |
| 47 | 2019/6/21 | L | Okayama |
| 48 | 2019/6/21 | L | Okayama |

| | | | |
|---|---|---|---|
| 49 | 2019/6/21 | L | Okayama |
| 50 | 2019/6/21 | M | Osaka |
| 51 | 2019/6/21 | M | Osaka |
| 52 | 2019/6/21 | M | Osaka |
| 53 | 2019/6/21 | M | Osaka |
| 54 | 2019/6/21 | M | Osaka |
| 55 | 2019/6/21 | D | Hiroshima |
| 56 | 2019/6/21 | D | Hiroshima |
| 57 | 2019/6/21 | N | Osaka |
| 58 | 2019/6/21 | N | Osaka |
| 59 | 2019/6/21 | N | Osaka |
| 60 | 2019/6/21 | N | Osaka |
| 61 | 2019/6/24 | A | Nara |
| 62 | 2019/6/24 | A | Nara |
| 63 | 2019/6/24 | O | Osaka |
| 64 | 2019/6/24 | O | Osaka |
| 65 | 2019/6/24 | C | Hyogo |
| 66 | 2019/6/24 | C | Hyogo |
| 67 | 2019/6/24 | I | Hyogo |
| 68 | 2019/6/24 | I | Hyogo |
| 69 | 2019/6/24 | N | Osaka |
| 70 | 2019/6/24 | N | Osaka |
| 71 | 2019/6/24 | P | Shiga |
| 72 | 2019/6/24 | P | Shiga |
| 73 | 2019/6/25 | Q | Okayama |
| 74 | 2019/6/25 | Q | Okayama |
| 75 | 2019/6/25 | Q | Okayama |
| 76 | 2019/6/25 | Q | Okayama |
| 77 | 2019/6/25 | Q | Okayama |
| 78 | 2019/6/25 | R | Okayama |
| 79 | 2019/6/25 | R | Okayama |
| 80 | 2019/6/25 | R | Okayama |
| 81 | 2019/6/25 | R | Okayama |
| 82 | 2019/6/25 | S | Osaka |
| 83 | 2019/6/25 | S | Osaka |
| 84 | 2019/6/25 | S | Osaka |
| 85 | 2019/6/25 | S | Osaka |
| 86 | 2019/7/16 | T | Wakayama |
| 87 | 2019/7/16 | T | Wakayama |
| 88 | 2019/7/16 | T | Wakayama |
| 89 | 2019/7/16 | T | Wakayama |
| 90 | 2019/7/16 | T | Wakayama |
| 91 | 2019/7/16 | U | Wakayama |
| 92 | 2019/7/16 | C | Hyogo |
| 93 | 2019/7/16 | C | Hyogo |
| 94 | 2019/7/16 | C | Hyogo |
| 95 | 2019/7/16 | E | Wakayama |
| 96 | 2019/7/16 | E | Wakayama |
| 97 | 2019/7/16 | F | Mie |
| 98 | 2019/7/16 | F | Mie |

| | | | |
|---|---|---|---|
| 99 | 2019/7/16 | F | Mie |
| 100 | 2019/7/16 | F | Mie |
| 101 | 2019/7/18 | G | Hiroshima |
| 102 | 2019/7/18 | G | Hiroshima |
| 103 | 2019/7/18 | G | Hiroshima |
| 104 | 2019/7/18 | G | Hiroshima |
| 105 | 2019/7/18 | G | Hiroshima |
| 106 | 2019/7/18 | I | Hyogo |
| 107 | 2019/7/18 | I | Hyogo |
| 108 | 2019/7/18 | I | Hyogo |
| 109 | 2019/7/18 | I | Hyogo |
| 110 | 2019/7/18 | I | Hyogo |
| 111 | 2019/7/18 | I | Hyogo |
| 112 | 2019/7/18 | I | Hyogo |
| 113 | 2019/7/18 | K | Nara |
| 114 | 2019/7/18 | K | Nara |
| 115 | 2019/7/18 | K | Nara |
| 116 | 2019/7/18 | K | Nara |
| 117 | 2019/7/19 | V | Nara |
| 118 | 2019/7/19 | V | Nara |
| 119 | 2019/7/19 | M | Osaka |
| 120 | 2019/7/19 | M | Osaka |
| 121 | 2019/7/19 | M | Osaka |
| 122 | 2019/7/19 | M | Osaka |
| 123 | 2019/7/19 | M | Osaka |
| 124 | 2019/7/19 | D | Hiroshima |
| 125 | 2019/7/19 | D | Hiroshima |
| 126 | 2019/7/19 | D | Hiroshima |
| 127 | 2019/7/19 | D | Hiroshima |
| 128 | 2019/7/19 | D | Hiroshima |
| 129 | 2019/7/19 | N | Osaka |
| 130 | 2019/7/19 | N | Osaka |
| 131 | 2019/7/19 | N | Osaka |
| 132 | 2019/7/24 | W | Kyoto |
| 133 | 2019/7/24 | W | Kyoto |
| 134 | 2019/7/24 | W | Kyoto |
| 135 | 2019/7/24 | W | Kyoto |
| 136 | 2019/7/25 | X | Shiga |
| 137 | 2019/7/25 | X | Shiga |
| 138 | 2019/7/25 | X | Shiga |
| 139 | 2019/7/25 | X | Shiga |
| 140 | 2019/7/25 | X | Shiga |
| 141 | 2019/7/25 | Y | Hyogo |
| 142 | 2019/7/25 | Y | Hyogo |
| 143 | 2019/7/25 | Y | Hyogo |
| 144 | 2019/7/25 | Y | Hyogo |
| 145 | 2019/7/25 | Y | Hyogo |
| 146 | 2019/7/26 | Z | Shimane |
| 147 | 2019/7/26 | Z | Shimane |
| 148 | 2019/7/26 | Z | Shimane |

| | | | |
|---|---|---|---|
| 149 | 2019/7/26 | Z | Shimane |
| 150 | 2019/7/26 | Z | Shimane |
| 151 | 2019/7/31 | AA | Kyoto |
| 152 | 2019/7/31 | AA | Kyoto |
| 153 | 2019/7/31 | AA | Kyoto |
| 154 | 2019/7/31 | AB | Gifu |
| 155 | 2019/7/31 | AB | Gifu |
| 156 | 2019/7/31 | AB | Gifu |
| 157 | 2019/7/31 | AB | Gifu |
| 158 | 2019/7/31 | AB | Gifu |
| 159 | 2019/7/31 | AB | Gifu |
| 160 | 2019/7/31 | AB | Gifu |
| 161 | 2019/8/2 | M | Osaka |
| 162 | 2019/8/2 | M | Osaka |
| 163 | 2019/8/2 | M | Osaka |
| 164 | 2019/8/2 | D | Hiroshima |
| 165 | 2019/8/2 | D | Hiroshima |
| 166 | 2019/8/2 | D | Hiroshima |
| 167 | 2019/8/2 | D | Hiroshima |
| 168 | 2019/8/2 | N | Osaka |
| 169 | 2019/8/2 | N | Osaka |
| 170 | 2019/8/2 | N | Osaka |
| 171 | 2019/8/7 | A | Nara |
| 172 | 2019/8/7 | A | Nara |
| 173 | 2019/8/7 | A | Nara |
| 174 | 2019/8/7 | A | Nara |
| 175 | 2019/8/7 | AC | Wakayama |
| 176 | 2019/8/7 | AC | Wakayama |
| 177 | 2019/8/7 | AC | Wakayama |
| 178 | 2019/8/7 | AC | Wakayama |
| 179 | 2019/8/7 | AC | Wakayama |
| 180 | 2019/8/7 | AC | Wakayama |
| 181 | 2019/8/7 | G | Hiroshima |
| 182 | 2019/8/7 | G | Hiroshima |
| 183 | 2019/8/7 | G | Hiroshima |
| 184 | 2019/8/7 | G | Hiroshima |
| 185 | 2019/8/7 | G | Hiroshima |
| 186 | 2019/8/7 | AD | Aichi |
| 187 | 2019/8/7 | AD | Aichi |
| 188 | 2019/8/7 | AD | Aichi |
| 189 | 2019/8/7 | AD | Aichi |
| 190 | 2019/8/7 | AD | Aichi |
| 191 | 2019/8/26 | AE | Kyoto |
| 192 | 2019/8/26 | AE | Kyoto |
| 193 | 2019/8/26 | AF | Wakayama |
| 194 | 2019/8/26 | AF | Wakayama |
| 195 | 2019/8/26 | AF | Wakayama |
| 196 | 2019/8/28 | X | Shiga |
| 197 | 2019/8/28 | X | Shiga |
| 198 | 2019/8/28 | X | Shiga |

| | | | |
|---|---|---|---|
| 199 | 2019/8/28 | X | Shiga |
| 200 | 2019/8/28 | X | Shiga |

Supporting Table 2. Bovine sera used in this study

| No. | Collection date | Vendor | Prefecture |
|---|---|---|---|
| 201 | 2019/10/8 | a | Kagoshima |
| 202 | 2019/10/8 | a | Kagoshima |
| 203 | 2019/10/8 | a | Kagoshima |
| 204 | 2019/10/8 | b | Wakayama |
| 205 | 2019/10/8 | b | Wakayama |
| 206 | 2019/10/8 | c | Hyogo |
| 207 | 2019/10/8 | c | Hyogo |
| 208 | 2019/10/8 | c | Hyogo |
| 209 | 2019/10/8 | c | Tokushima |
| 210 | 2019/10/8 | c | Tokushima |
| 211 | 2019/10/8 | c | Tokushima |
| 212 | 2019/10/8 | d | Oita |
| 213 | 2019/10/8 | d | Oita |
| 214 | 2019/10/8 | d | Oita |
| 215 | 2019/10/8 | d | Oita |
| 216 | 2019/10/8 | e | Hokkaido |
| 217 | 2019/10/9 | f | Shimane |
| 218 | 2019/10/9 | f | Shimane |
| 219 | 2019/10/9 | f | Shimane |
| 220 | 2019/10/9 | g | Tokushima |
| 221 | 2019/10/9 | g | Tokushima |
| 222 | 2019/10/9 | h | Miyazaki |
| 223 | 2019/10/9 | h | Miyazaki |
| 224 | 2019/10/9 | h | Miyazaki |
| 225 | 2019/10/9 | i | Saga |
| 226 | 2019/10/9 | i | Saga |
| 227 | 2019/10/9 | i | Saga |
| 228 | 2019/10/9 | k | Shizuoka |
| 229 | 2019/10/9 | k | Shizuoka |
| 230 | 2019/10/9 | j | Aichi |
| 231 | 2019/10/9 | k | Shizuoka |
| 232 | 2019/10/9 | k | Shizuoka |
| 233 | 2019/10/17 | l | Yamaguchi |
| 234 | 2019/10/17 | l | Yamaguchi |
| 235 | 2019/10/17 | l | Yamaguchi |
| 236 | 2019/10/17 | m | Hyogo |
| 237 | 2019/10/17 | n | Okayama |
| 238 | 2019/10/17 | n | Okayama |
| 239 | 2019/10/17 | o | Hiroshima |
| 240 | 2019/10/17 | o | Hiroshima |
| 241 | 2019/10/17 | o | Hiroshima |
| 242 | 2019/10/21 | p | Ehime |
| 243 | 2019/10/21 | p | Hiroshima |
| 244 | 2019/10/21 | p | Hiroshima |
| 245 | 2019/10/21 | p | Hiroshima |
| 246 | 2019/10/21 | p | Okayama |
| 247 | 2019/10/21 | p | Okayama |
| 248 | 2019/10/21 | q | Wakayama |

| | | | |
|---|---|---|---|
| 249 | 2019/10/21 | q | Wakayama |
| 250 | 2019/10/21 | q | Wakayama |
| 251 | 2020/3/10 | p | Hiroshima |
| 252 | 2020/3/10 | p | Hiroshima |
| 253 | 2020/3/10 | p | Hiroshima |
| 254 | 2020/3/10 | c | Tokushima |
| 255 | 2020/3/10 | c | Tokushima |
| 256 | 2020/3/10 | c | Tokushima |
| 257 | 2020/3/10 | d | Oita |
| 258 | 2020/3/10 | d | Oita |
| 259 | 2020/3/10 | d | Oita |
| 260 | 2020/3/12 | r | Saga |
| 261 | 2020/3/12 | r | Saga |
| 262 | 2020/3/12 | r | Saga |
| 263 | 2020/3/12 | s | Tokushima |
| 264 | 2020/3/12 | s | Tokushima |
| 265 | 2020/3/12 | s | Tokushima |
| 266 | 2020/3/12 | n | Okayama |
| 267 | 2020/3/12 | n | Okayama |
| 268 | 2020/3/12 | n | Okayama |
| 269 | 2020/3/12 | j | Aichi |
| 270 | 2020/3/12 | j | Aichi |
| 271 | 2020/3/12 | j | Aichi |
| 272 | 2020/3/13 | t | Nagasaki |
| 273 | 2020/3/13 | t | Nagasaki |
| 274 | 2020/3/13 | t | Nagasaki |
| 275 | 2020/3/13 | t | Nagasaki |
| 276 | 2020/3/13 | t | Nagasaki |
| 277 | 2020/3/13 | t | Nagasaki |
| 278 | 2020/3/17 | u | Nagasaki |
| 279 | 2020/3/17 | u | Nagasaki |
| 280 | 2020/3/17 | v | Hokkaido |
| 281 | 2020/3/17 | v | Hokkaido |
| 282 | 2020/3/17 | v | Hokkaido |
| 283 | 2020/3/17 | w | Tottori |
| 284 | 2020/3/17 | w | Tottori |
| 285 | 2020/3/17 | w | Tottori |
| 286 | 2020/3/17 | x | Ehime |
| 287 | 2020/3/17 | x | Ehime |
| 288 | 2020/3/17 | x | Ehime |
| 289 | 2020/3/23 | y | Okayama |
| 290 | 2020/3/23 | z | Hyogo |
| 291 | 2020/3/23 | z | Hyogo |
| 292 | 2020/3/23 | z | Hyogo |
| 293 | 2020/3/23 | 1 | Tokushima |
| 294 | 2020/3/23 | 2 | Nagano |
| 295 | 2020/3/23 | 2 | Nagano |
| 296 | 2020/3/23 | 2 | Nagano |
| 297 | 2020/3/23 | 3 | Wakayama |
| 298 | 2020/3/23 | i | Saga |

| | | | |
|---|---|---|---|
| 299 | 2020/3/23 | 4 | Kumamoto |
| 300 | 2020/3/23 | 4 | Kumamoto |
| 301 | 2020/3/23 | 4 | Kumamoto |
| 302 | 2020/3/23 | 4 | Kumamoto |
| 303 | 2020/3/23 | e | Miyazaki |
| 304 | 2020/3/23 | e | Miyazaki |
| 305 | 2020/3/23 | e | Miyazaki |

Supporting Table 2. Primers for cloning

| Name | Sequence (5' – 3') |
|---|---|
| faeG1-Fw | TAGTCGACAAGTTGCCATGGTGTCCGGCTC |
| faeG1-Rv | ACGCGGCCGCTTATTGATAAGAAACAACGA |
| faeG2-Fw | TAGTCGACAAATGGTCTCCACACCTGCGCT |
| faeG2-Rv | ACGCGGCCGCTTAATTATAAGTTACTGCAA |
